# Summary tests of introgression are highly sensitive to rate variation across lineages

**DOI:** 10.1101/2023.01.26.525396

**Authors:** Lauren E. Frankel, Cécile Ané

## Abstract

The evolutionary implications and frequency of hybridization and introgression are increasingly being recognized across the tree of life. To detect hybridization from multi-locus and genome-wide sequence data, a popular class of methods are based on summary statistics from subsets of 3 or 4 taxa. However, these methods often carry the assumption of a constant substitution rate across lineages and genes, which is commonly broken in many groups. In this work, we quantify the effects of rate variation on the *D*-statistic (also known as ABBA-BABA test), the *D*_3_ statistic, and HyDe. All three tests are used widely across a range of taxonomic groups, in part because they are very fast to compute. We consider rate variation across species lineages, across genes, their lineage-by-gene interaction, and rate variation across gene-tree edges. We simulated species networks according to a birth-death-hybridization process so as to capture a range of realistic species phylogenies. For all three methods tested, we found a marked increase in the false discovery of reticulation (type-1 error rate) when there is rate variation across species lineages. The *D*_3_ statistic was the most sensitive, with around 80% type-1 error, such that *D*_3_ appears to more sensitive to a departure from the clock than to the presence of reticulation. For all three tests, the power to detect hybridization events decreased as the number of hybridization events increased, indicating that multiple hybridization events can “hide” one another if they occur within a small subset of taxa. Our study highlights the need to consider rate variation when using site-based summary statistics, and points to the advantages of methods that do not require assumptions on evolutionary rates across lineages or across genes.

## 1 Introduction

Hybridization, the exchange of genes between distinct evolutionary lineages and subsequent backcrossing is a well-documented evolutionary phenomenon in animals, plants, and fungi. Particularly in plants, it can play an important role in speciation and adaptive evolution [Edelman and Mallet, 2021, Soltis and Soltis, 2009]. Interspecific hybrids have been documented in 40% of vascular plant families and in 16% of vascular plant genera [Whitney et al., 2010]. As sequencing abilities progressed at the turn of the century, so have phylogenetic methods to detect and model hybridization, gene flow and migration between populations. There are two main classes of hybridization detection methods. Faster methods aim to test for the presence of reticulation. Most of these methods are based on summary statistics calculated from site and gene tree frequencies on small subsets of (3-5) taxa, such as the *D*-statistic [Green et al., 2010, Durand et al., 2011], *D*_3_ [Hahn and Hibbins, 2019], HyDe [Blischak et al., 2018], QuIBL [Edelman et al., 2019], MSCquartets [Rhodes et al., 2021], and the *f*_3_ statistic [Patterson et al., 2012]. In the second class, model-based likelihood methods aim to estimate a phylogenetic network on the full taxon set. These methods are more computationally intensive. The most widely-used of these methods are those that scale to larger taxon sets such as such as SNaQ [Solís-Lemus and Ané, 2016], PhyloNet-MPL [Yu and Nakhleh, 2015] or TreeMix [Pickrell and Pritchard, 2012]. By estimating a species network, these methods gain insights into the timing, direction, and magnitude of hybridization events.

While these methods hold great promise in identifying reticulate and admixture events, much is still unknown about their comparative performances when model assumptions are violated. In particular, one common model assumption is a single substitution rate across all genes and lineages. Substitution rates are known to vary depending on gene and lineage, especially beyond shallow time scales [Gaut et al., 2011, Baer et al., 2007]. Cao et al. [2022] found that the Bayesian species network inference method `MCMC-SEQ` [Wen and Nakhleh, 2018], implemented in `PhyloNet`, is sensitive to substitution rate variation across genes, resulting in the estimation of extra reticulation events in both empirical and simulated datasets. Consequently, Cao et al. [2022] added support for per-locus rate variation in `MCMC-SEQ`. Yet, the robustness of various hybridization detection methods to rate variation has not yet been assessed and quantified. In this paper, we do so for the detection methods *D, D*_3_, and HyDe.

The *D*-statistic, also known as Patterson’s *D* or the ABBA-BABA test, is widely used [Green et al., 2010, Durand et al., 2011]. It is based on biallelic site pattern counts for groupings of four taxa. These four taxa should have a known asymmetric tree under the null hypothesis of no reticulation, which can be represented as (((*t*_1_, *t*_2_), *t*_3_), *t*_4_) where *t*_4_ is the outgroup taxon. It is defined as

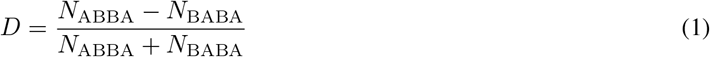

where *N*_ABBA_ is the number of sites for which taxa *t*_2_ and *t*_3_ have a base B and taxa *t*_1_ and *t*_4_ have a base A, different from B. Similarly, *N*_BABA_ is the number of sites for which taxa *t*_1_ and *t*_3_ share base B and taxa *t*_2_ and *t*_4_ share a different base A. It is typically assumed that base A is ancestral and B is derived, but this assumption is not checked or necessary. If biallelic site patterns ABBA and BABA occur in roughly equal proportions, this is assumed to be the signature of incomplete lineage sorting (ILS) only. However, if one pattern occurs in significantly greater frequency than the other (assessed with bootstrap or jackknife resampling), then directional hybridization between *t*_3_ and either of *t*_1_ or *t*_2_ is inferred.

This test of reticulation assumes a clock, although this assumption was not explicit in its original presentation, which was applied at a shallow evolutionary scale (human evolution). The ABBA-BABA test was shown to be sensitive to a violation of the clock in a small simulation study [Blair and Ané, 2020]. Other assumptions of the *D*-statistic include infinite sites (no saturation of bases), no ancestral population structure, a single historical introgression event under the alternative hypothesis, no introgression between the outgroup and ingroups, and that all introgressed taxa are sampled. This last assumption has been the focus of a recent study where “ghost” lineages (unsampled or extinct populations) can result in incorrect interpretations of donors and recipients of gene flow [Tricou et al., 2022a].

HyDe, a relative of the *D*-statistic, is notable in that it aims to identify the putative parent and hybrid populations within a set of three ingroup taxa and an outgroup, assuming hybrid speciation, that is, lineage generative hybridization [Blischak et al., 2018]. By testing each taxon in a triplet as the possible hybrid, it generates three statistics per triplet of ingroup taxa. HyDe also estimates the proportion of genomic contribution from a parent species to the hybrid species, *γ*. The test statistic in HyDe uses the ratio between *f*_1_ = *N*_ABBA_ − *N*_BABA_ and *f*_2_ = *N*_BBAA_ − *N*_BABA_. Like the *D* test, it assumes a molecular clock. It also assumes that sites are unlinked, that is, sites have independent gene trees. In simulation studies with a variety of hybridization scenarios, HyDe performed well with low false discovery rate, high power to detect hybridization and accurate estimation of *γ*, except under high ILS [Kong and Kubatko, 2021] and when hybridization occurred in deep time or appeared to involved a ghost lineage [Bjørner et al., 2022].

*D*_3_ is another relative of the *D*-statistic, but uses pairwise genetic distances instead of 4-taxon biallelic site patterns, and does not require an outgroup [Hahn and Hibbins, 2019]. For three ingroup taxa with known tree ((*t*_1_, *t*_2_), *t*_3_) under the null hypothesis of no reticulation, *D*_3_ is defined as

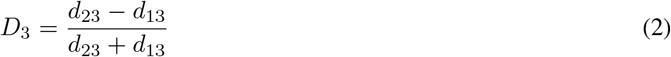

where *d*_13_ and *d*_23_ are the pairwise genetic distance between each sister taxon (*t*_1_ or *t*_2_) and *t*_3_, measured by the number of sites at which their sequences differ. For *D*_3_ to detect reticulation between non-sister taxa (*t*_3_ and either *t*_1_ or *t*_2_), Hahn and Hibbins [2019] assume an infinite sites model with a constant mutation rate, an ultrametric phylogeny (equal number of generations from the root to all sampled populations) and no ancestral population structure. Hahn and Hibbins [2019] find that the *D*_3_ statistic results in high rates of false positives above a ∼0.01% difference in substitution rates between taxa. Similarly to *D, D*_3_ assumes correct taxon sampling, as it may result in erroneous interpretations of introgressed taxa when ghost lineages are involved [Tricou et al., 2022b].

In this work, we aim to assess and quantify the effect of substitution rate variation on the accuracy of *D, D*_3_ and HyDe for detecting reticulation.

## 2 Methods

Rather than using a small number of fixed networks, we simulated species networks under a birth-death-hybridization process, and then simulated DNA alignments under these varied networks. Gene trees were generated from the multispecies network coalescent model, and their edge lengths were converted to units in substitutions per site after generating various modes of rate variation, as described below.

### 2.1 Simulating networks, gene trees, and sequences

#### 2.1.1 Network simulation

We simulated networks, each with 10 taxa, under the birth-death-hybridization process [Justison et al., 2022b] with the `R` package `SiPhyNetwork` v.1.0.0 [Justison et al., 2022a]. We used a speciation rate *λ* = 1, an extinction rate of *μ* = 0.2, and a reticulation rate of *ν* = 0.3 reticulations (per pair of lineages). The proportion of reticulations that were lineage generative (hybrid speciation), lineage degenerative (fusion), and lineage neutral (gene flow) were set to 0.5, 0.25, and 0.25 respectively. For each proposed reticulation event, the probability of success was a stepwise function of the genetic distance between the two proposed reticulating lineages, with a single step occurring at 0.75. In other words, any proposed reticulation between lineages of average genetic distance *d* was rejected if *d >* 0.75 and accepted if *d* ≤ 0.75. Inheritance probabilities of hybrid edges were drawn from a beta distribution with shape parameters 2 and 2, whose mean is 0.5 and standard deviation is 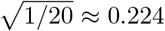. We then shrunk any 2-cycle and 3-cycle iteratively, because reticulations in such cycles are between sister species and are not detectable by many methods, including the tests considered here. A 2-cycle corresponds to two parallel hybrid edges that share the same parent and child populations. It represents a population splitting into 2 separate populations, which then fuse back into a single population. Shrinking this 2-cycle means replacing the two parallel hybrid edges by a single tree edge. Shrinking a 3-cycle means replacing a triangle including 2 partner hybrid edges by a 3-edge subtree, while preserving the average distance between the 3 nodes in the triangle. In each simulated network, the ingroup and outgroup taxa were defined as the groups of taxa descendant from each of the two edges adjacent to the root, with the outgroup being the smallest of the two groups. Finally, we rejected any network that did not meet the following criteria: 10 species total (using the simple sampling approach), no reticulation between ingroup and outgroup clades, and between 1 and 5 reticulation events. The ingroup necessarily had 5 or more taxa, since it was defined as being no smaller than the outgroup. It was possible for a network to have no reticulation within the ingroup clade, in which case reticulations were restricted to the outgroup clade and the ingroup phylogeny satisfied the null hypothesis of no reticulation.

Once we had a network that met our filters, we scaled all its branches to have a median edge length of 1.0, which we used as measuring coalescent units, to later simulate a moderate level of ILS, similar across all simulated networks.

We calculated a scaler *L* for each network, for later conversion of coalescent units into substitutions per site, to obtain a level of sequence variability within the ingroup clade that was similar across all networks. Namely, we extracted the major tree from each species network and recorded the length *L* (sum of all edge lengths) of the ingroup clade in this major tree, measured in coalescent units.

#### 2.1.2 Gene tree simulation

From each scaled network, we simulated 1000 gene trees under the network multispecies coalescent (NMSC) using `PhyloCoalSimulations` v0.1.0 [Fogg et al., 2023, Fogg and Ané, 2022], written in `Julia` [Bezanson et al., 2017]. These simulated gene trees had edge lengths in coalescent units, and had degree-2 nodes to mark each speciation and reticulation event (node in the species network, see Fig. 2). Thanks to these extra nodes, each edge in a gene tree mapped to a unique population lineage in the species network.

To convert edge lengths to a number of expected substitutions per site, we multiplied all edge lengths in all gene trees by 0.03*/L* substitutions per site per coalescent unit, where *L* was network-specific as described above. This scaling factor lead to an expected 0.03 substitutions/site in the ingroup phylogeny. For alignments with 1000 sites per locus, this choice provided enough sequence variation to perform site-based statistical analyses later without saturating sequences.

#### 2.1.3 Simulation of substitution rate variation

To simulate variation in the rate of molecular evolution, we considered four different levels influencing substitution rates: at the species (or lineage) level, at the gene level, their interaction, and at the level of individual edges within each gene trees. More specifically, for each edge *e* in each gene tree — augmented by degree-2 nodes at times crossing speciation and reticulation events — the edge length *𝓁*(*e*) in substitutions per site was determined by the gene *g* and the species lineage *l* that the edge evolved in, as follows:

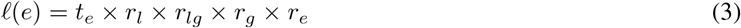

where rate multipliers are described below, and *t*_*e*_ is the length of *e* after simulation under the coalescent and after conversion of coalescent units to expected substitutions/site, as described in the previous section. Note that each gene tree is ultrametric when edge lengths *t*_*e*_ are used: all the paths from the root to any leaf have the same length when using *t*_*e*_, prior to applying rate multipliers with (3). Also, since we mapped each gene tree to the species network, each edge *e* corresponds to what may otherwise be considered an ‘edge segment’ if degree-2 nodes were not considered in gene trees.

Each lineage *l* in the species network was assigned a relative rate *r*_*l*_, that affected all edges from this lineage across all genes. We simulated *r*_*l*_ independently across species lineages, from a log-normal distribution with mean 1 and standard deviation 0 or 0.7 on the log scale, that is, the standard deviation of log(*r*_*l*_) was either set to 0 (no rate variation) or to 0.7. This value was chosen to generate sufficient substitution rate variation (Fig. 1).

**Figure 1:**
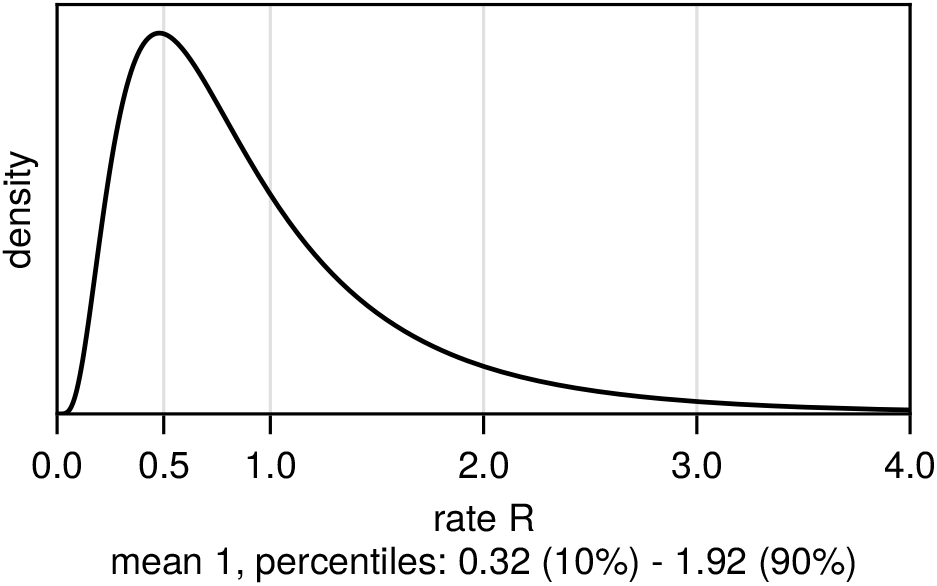
Density of the log-normal distribution used for simulating rates *R*, with standard deviation of 0.7 on the log-scale. To set the mean of *R* to 1, the mean of log(*R*) was set to −0.72^2^*/*2.

For each combination of lineage *l* and gene *g*, we also simulated a lineage-by-gene specific rate *r*_*lg*_, which affected all edges of a given gene that evolved in the same species lineage. Sometimes *r*_*lg*_ affected a single edge only, and sometimes multiple edges (Fig. 2). Rates *r*_*lg*_ were simulated independently across genes and lineages, from a log-normal distribution with mean 1, and with standard deviation of 0 (rate homogeneity) or 0.7 as for lineage-specific rates (Fig. 1).

**Figure 2:**
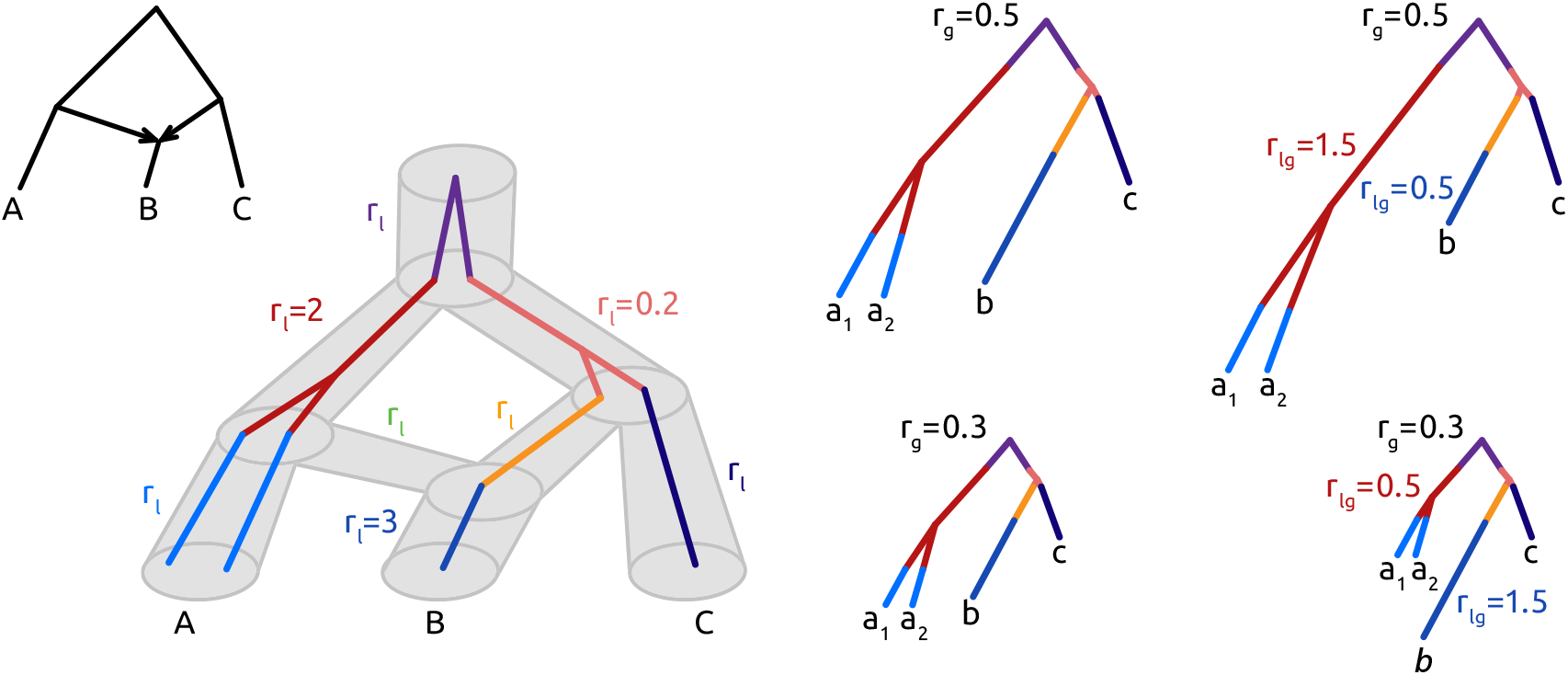
Illustration of the different types of rate variation: across lineages, across genes, and lineage-by-gene specific. Left: example network, first drawn with thin black edges (leftmost), then drawn with gray cylinders (middle-left) within which a gene tree is simulated from the coalescent with two individuals sampled from population A. The same gene tree topology is shown on the right, with its edges color-coded by the species lineage in which it evolved. Each species lineage in the network has a lineage-specific rate *r*_*l*_, set to 1 unless indicated. For example, *r*_*l*_ = 3 for species B indicates that dark blue gene edges, which evolved in the species B lineage, evolved three times as fast as expected on average. Right: gene trees with edges drawn proportionally to their lengths in substitutions/site from (3) with a gene-specific rate *r*_*g*_ = 0.5 (top) or *r*_*g*_ = 0.3 (bottom), and the addition of lineage-by-gene specific rates *r*_*lg*_ (rightmost). Multiple edges of the same gene may evolve in the same lineage, and therefore share the same rate multiplier *r*_*lg*_. This illustration assumes *no* residual edge variation (*r*_*e*_ = 1 for all edges).

The gene-specific rate *r*_*g*_ was used to affect all edges in the same gene tree, for the simulation of fast-evolving genes and slow-evolving genes. Gene-specific rates were drawn independently across genes, from a log-normal distribution as before: with mean 1 and standard deviation 0 (homogeneous rates) or 0.7 (rate variation, Fig. 1). Finally, a residual edge rate for edge *e* in gene *g, r*_*e*_, independent of the edge’s lineage, was also drawn from a log-normal distribution with mean 1 and standard deviation 0 or 0.7. Combining these rate multipliers together leads to (3). With these four types of rate variation, there were 16 combinations of parameters. For each parameter set, we simulated 100 independent replicates, that is, 100 networks and 1000 genes along each network.

After obtaining gene trees with edge lengths scaled to substitutions per site (with possible rate variation), sequences were simulated along gene trees with `Seq-Gen` v.1.3.4 [Rambaut and Grass, 1997] under the HKY model (different rate of transitions and transversions as well as unequal base frequencies of the four nucleotides). Starting base frequencies were 0.2 A, 0.3 C, 0.3 G and 0.2 T, with a transition/transversion ratio of 3. We also simulated rate variation across sites, under a discretized gamma distribution with shape *α* = 0.35 and 10 rate categories for the discretization. We wrote sequence alignments in linear fasta format, and converted to phylip format for use in HyDe using the `fasta2phylip.py` script [Chafin, 2019, Chafin et al., 2021].

### 2.2 Introgression inference

To calculate the *D*-statistic (1), we used the Julia implementation from Blair and Ané [2020], expanded to perform locus-resampling bootstrap. We calculated the *D*-statistic for each possible ingroup triplet and a single outgroup taxon for each locus. If there were more than one outgroup taxon, one of them was chosen at random and the remainder were excluded from analyses. The same outgroup taxon was chosen all ingroup triplets. To reflect real data analysis where the topology of a triplet is unknown, we summed pattern counts for the three possible biallelic patterns where two taxa each have a different allele (ABBA, BABA, or BBAA). We then took the two smallest values as ABBA and BABA, and we took the most frequent pattern as reflecting the majority tree (BBAA). If the ABBA and BABA counts were both zero across all loci for a given ingroup triplet, then the *D* value, standard error, z value, and p value were considered missing for that triplet. Otherwise, the standard error of the *D* statistic, SE(*D*), was estimated as the standard deviation of *D* across 5000 bootstrap samples, resampling loci. By resampling loci, the *D* test accounts for linked SNPs within each locus. The *z*-value was calculated as *z* = *D/*SE(*D*). Finally, the two-tailed *p*-value was calculated from *z* and the cumulative probability density function of the normal distribution: *p* = 2IP*{Z >* |*z*|*}*.

For HyDe, we used `run_hyde.py` v.0.4.3 [Blischak et al., 2018] to analyze all possible triplets with a single outgroup taxon. The same outgroup taxon was used as for the *D* test above. We calculated the HyDe statistic for each of the 3 possible choices of the putative hybrid species for a given ingroup triplet, and we only retained the choice with the smallest p-value. This procedure matches real data analysis, when we do not have knowledge of the true topology for the species in the triplet. We applied a Bonferroni correction to account for the 3 tests within each triplet, by multiplying the retained (smallest) p-value by 3.

We implemented the calculation of the *D*_3_ statistic (2) in Julia to analyze all sets of 3 ingroup taxa. Similarly to *D*, to emulate real data analysis, we did not use knowledge of the true triplet topology to ascertain which of the two taxa were sister relative to the third, under the null hypothesis of no reticulation. Instead we took pair {*i, j*} as the sister pair if its pairwise distance *d*_*ij*_ was the smallest of the three pairs. The largest two pairwise distances were then used as *d*_13_ and *d*_23_ to calculate *D*_3_ from (2). Again, if *d*_13_ and *d*_23_ were both zero within all loci for a given triplet, the *D*_3_ value, its standard error, z value, and p value were considered missing for that triplet. Otherwise, the standard error for *D*_3_, SE(*D*_3_), was estimated as its standard deviation across 5000 bootstrap samples, resampling loci to account for linked sites within a locus. As for *D*, we then calculated *z* = *D*_3_*/*SE(*D*_3_) and the two-tailed *p*-value using a normal approximation *p* = 2IP*{Z >* |*z*|*}*.

For each set of 3 ingroup taxa, we extracted the subnetwork induced by the 3 taxa and the chosen outgroup. We had filtered out any species networks with reticulations between the ingroup and outgroup, so all reticulations in the subnetwork were restricted to the 3 ingroup taxa, or to the branch leading to the outgroup representative (e.g. via 2-cycles). We then calculated the number *h* of reticulations within the ingroup subnetwork. If *h* was 0, then the null hypothesis was true for the *D, D*_3_ and HyDe tests. If the ingroup subnetwork had *h* ≥ 1 reticulations, then we considered the fact that some of these reticulations may be undetectable by the tests, such as gene flow between sister taxa. Therefore we calculated the number of reticulations *h*′ in the subnetwork after iteratively shrinking any 2-cycle and 3-cycle. This elimination of undetectable reticulations was performed on the full 10-taxon network prior to simulating gene trees and rate variation. We repeated the operation on subnetworks to get *h*′, because a reticulation between non-sister taxa in the full network may appear as a reticulation between sister taxa after sampling a subset of 3 ingroup taxa, and thus become undetectable on this small taxon subset (see examples in Fig. 3 (a-b) and (d-e)).

**Figure 3:**
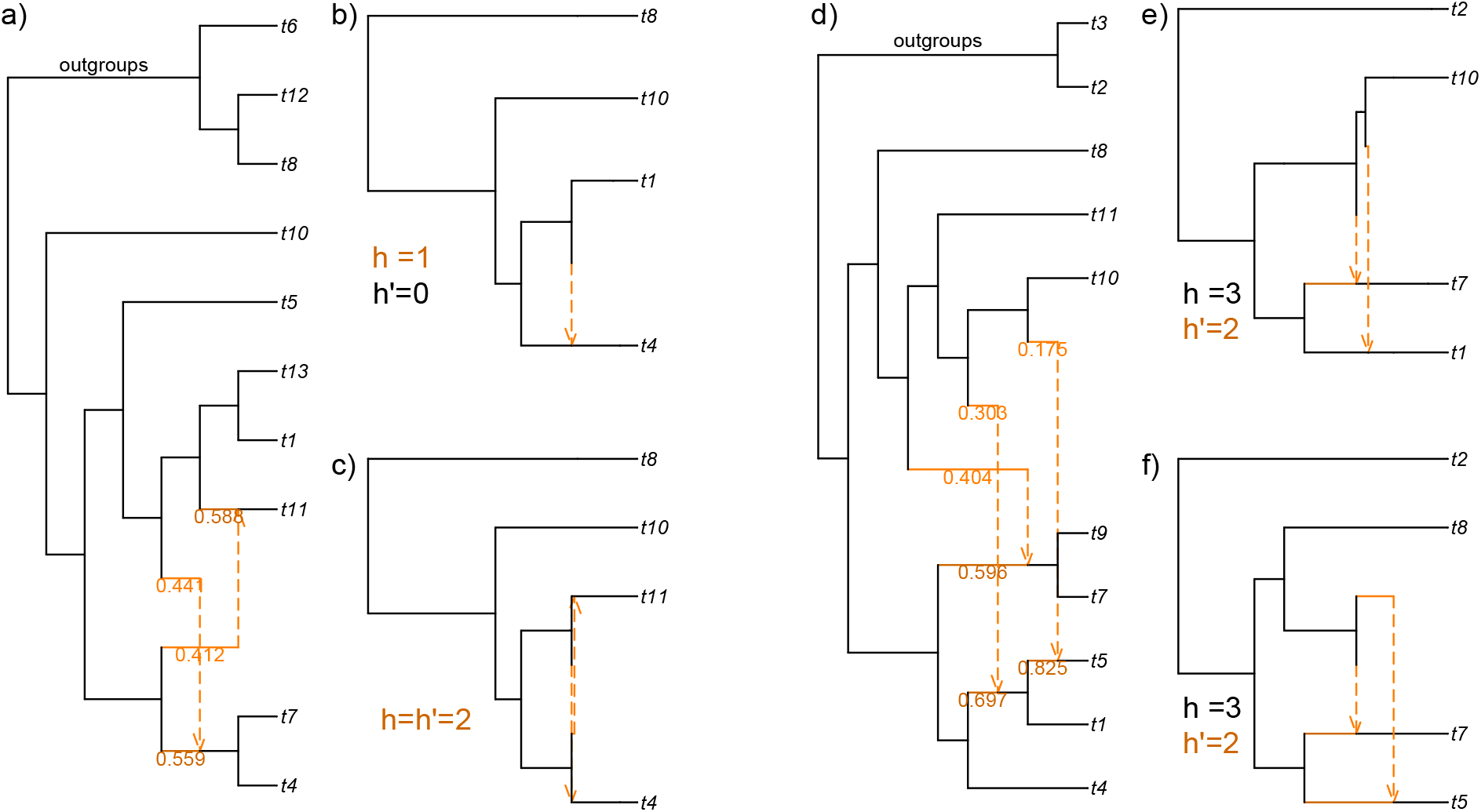
Example of simulated networks and subnetworks from 3 ingroup taxa and one outgroup taxon, for which reticulations go undetected by all methods in the absence of rate variation. (a) Simulated network on 10-taxon set. (b) Subnetwork from (a) with *h* = 1 yet *h*′ = 0 after shrinking 2- and 3-cycles. Lines are drawn proportional to edge lengths. (c) Subnetwork from (a) on a different set of 3 ingroup taxa, with *h* = *h*′ = 2 reticulations. (d) Simulated network on 10-taxon set from a different replicate. (e) and (f) Subnetworks from (d) with reticulations shown after shrinking 2/3-cycles (*h*′), in which reticulations “hide” each other.

### 2.3 Type-1 error versus power

A type-1 error is defined as rejecting the null hypothesis when it is true. For our analyses, the null hypothesis is that *h* = 0. But in fact neither *D, D*_3_ nor HyDe can detect reticulation between sister species. Thus, the null hypothesis should consider the network after removing reticulations between sister species, which are invisible to the tests considered here. This is leading us to a second definition of type-1 error rate: the probability of a significant test (as evaluated by a p-value lower than some threshold, *α* = 0.05 here) under the null hypothesis that *h*′ = 0, with *h*′ as defined above. Recall that *h*′ ≤ *h*, so *h*′ = 0 whenever *h* = 0, and this second definition of type-1 error extends the null hypothesis to more subnetworks.

To quantify the methods’ power, the probability of correctly rejecting the null hypothesis, we also considered two definitions, depending on *h* or *h*′: either when the alternative hypothesis is defined as “*h* ≥ 1” or as “*h*′ ≥ 1”.

## 3 Results

### 3.1 Type-1 error: false detection of reticulation

In the absence of rate variation and on taxon triplets whose phylogeny had no reticulation, all methods performed well, as measured by a rejection rate near 5% when comparing the test p-value to a significance threshold of *α* = 0.05 (orange filled circles with orange outline, on the left for each method, in Fig. 4 when *h* = 0 and Fig. 5 when *h*′ = 0). HyDe was above the 5% threshold, with a rejection rate of 9.6% when *h* = 0, and 7.4% when *h*′ = 0.

**Figure 4:**
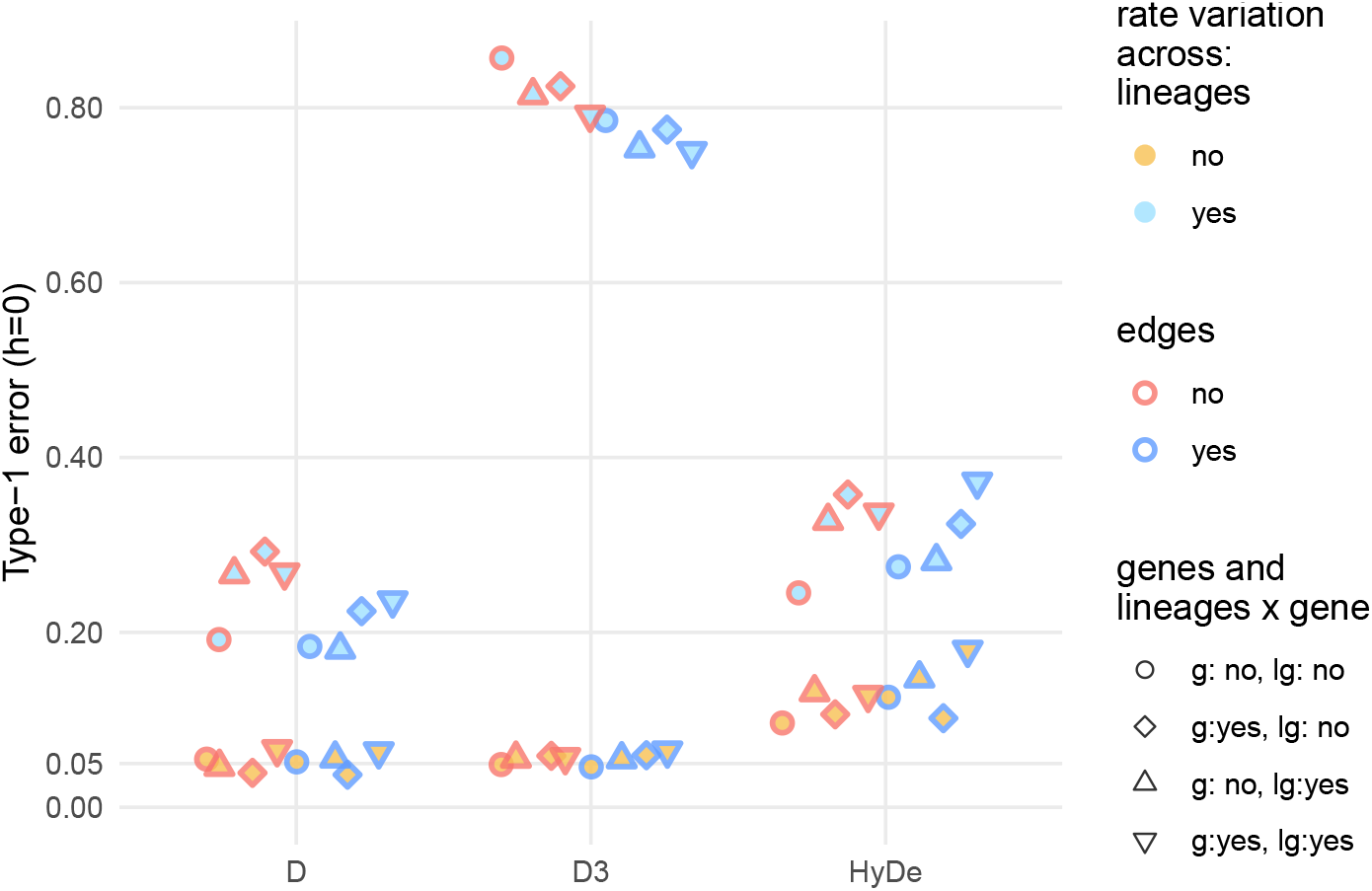
Type-1 error for *D, D*_3_, and HyDe when *h* = 0, that is, when there are no hybridization events before iteratively shrinking 2- and 3-cycles. Each point represents the proportion of tests with a p-value below 0.05 across four-taxon subnetworks from 100 simulated networks, with an average of 2232 four-taxon subnetworks per point. Blue-filled shapes denote the presence of lineage-rate variation (*r*_*l*_), and orange-filled shapes denote its absence. Blue outlines denote the presence of edge-rate variation (*r*_*e*_), and orange outlines denote its absence. Circles had neither gene-rate variation (*r*_*g*_), nor lineage-by-gene (*r*_*lg*_). Diamonds had gene-rate variation, but no lineage-by-gene. Upward pointing triangles did not have gene-rate variation, but did have lineage-by-gene. Downward pointing triangles had both gene and lineage-by-gene rate variation.

**Figure 5:**
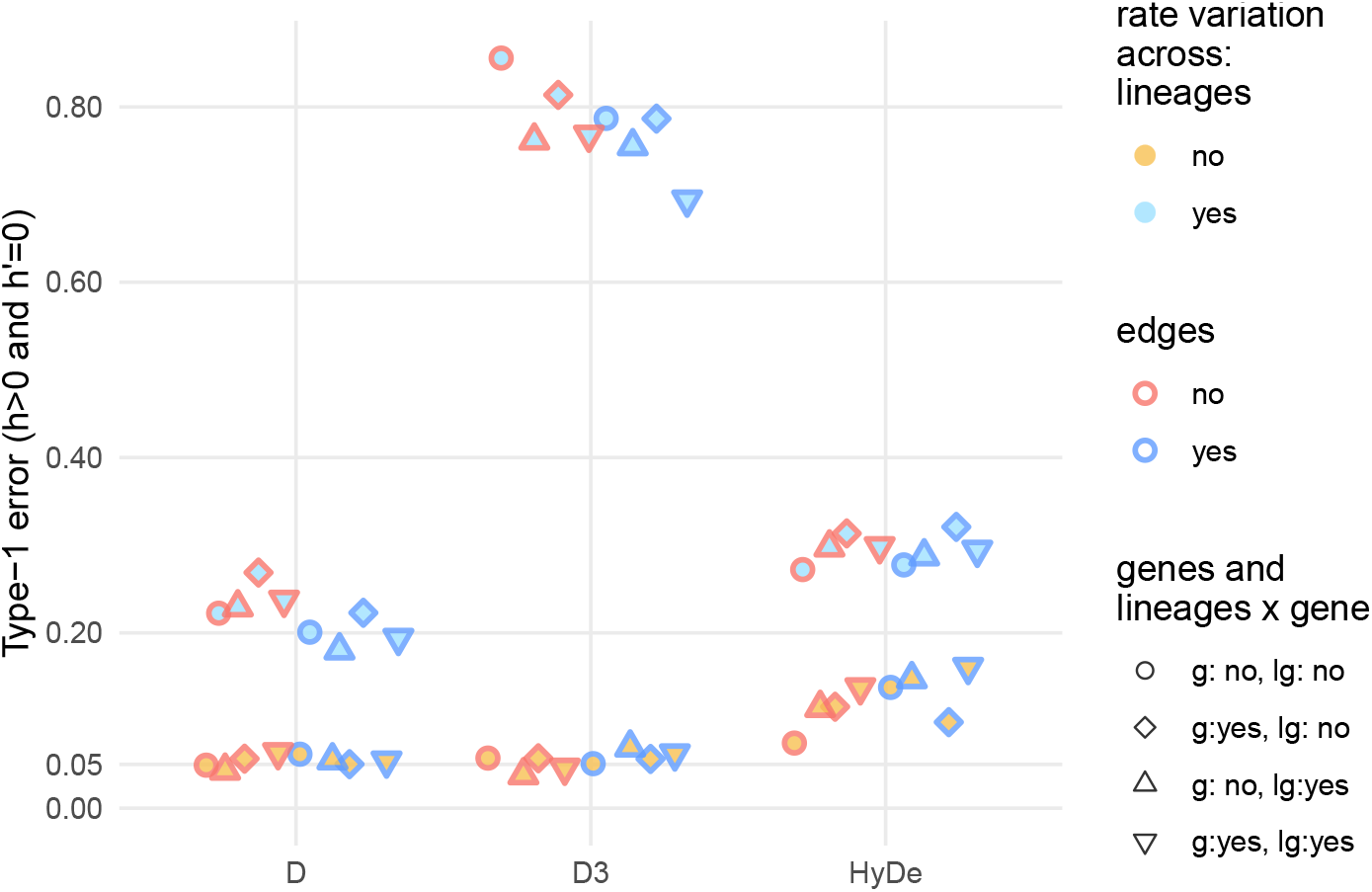
Type-1 error for *D, D*_3_, and HyDe when *h*′ ≥ 1 in the subnetwork prior to iteratively shrinking 2- and 3-cycles but all reticulations are eliminated (*h*′ = 0) after such shrinkage. Each point represents the proportion of tests with a p-value below 0.05 across four-taxon subnetworks from 100 simulated networks, with an average of 1864 four-taxon subnetworks per point. Fill color, outline color and point shape are as in Fig. 4.

Lineage rate variation (blue-filled points) had a large effect across all three tests and when either *h* or *h*′ = 0, with clusters of higher type-1 error rates when there was lineage rate variation, regardless of the presence or absence of other types of rate variation. With rate variation across species lineages, the type-1 error rate of *D* hovered around 20%, and that of HyDe hovered around 30% (Figs. 4 and 5). These rates would typically be considered as unacceptably high. The type-1 error rate of the *D*_3_ test was even higher, above 69% in all cases with lineage-rate variation, and with an average of 77.8% (across parameters for the other types of rate variation).

Other types of rate variation had some effect on type-1 error, but much smaller in magnitude than lineage-rate variation (Fig. 4). Rate variation across edges somewhat limited the adverse effect of lineage-rate variation for *D* and particularly *D*_3_. For HyDe, edge-rate variation somewhat increased type-1 error in the absence of lineage-rate variation (to about 15%).

### 3.2 Power to detect the presence of reticulations

When there remained hybridization events in the triplet subnetwork after shrinking 2/3-cycles, the power to reject the no-reticulation hypothesis varied upon the test and the number of hybridizations in the subnetwork. When *h*′ = 1 (Fig. 6), the power of the *D*-test is high between 60-80% (depending on the presence or absence of the various types of rate variation). HyDe’s power was higher, around or above 80%. For *D* and HyDe, there was no discernible pattern between power and the type of rate variation (Fig. 6). In contrast, *D*_3_’s power to detect reticulation was highly impacted by lineage-rate variation: its power was lower (around 60%) in the absence of lineage-rate variation (orange-filled points in Fig. 6), and higher (most often above 80%) in the presence of lineage-rate variation (blue-filled points).

**Figure 6:**
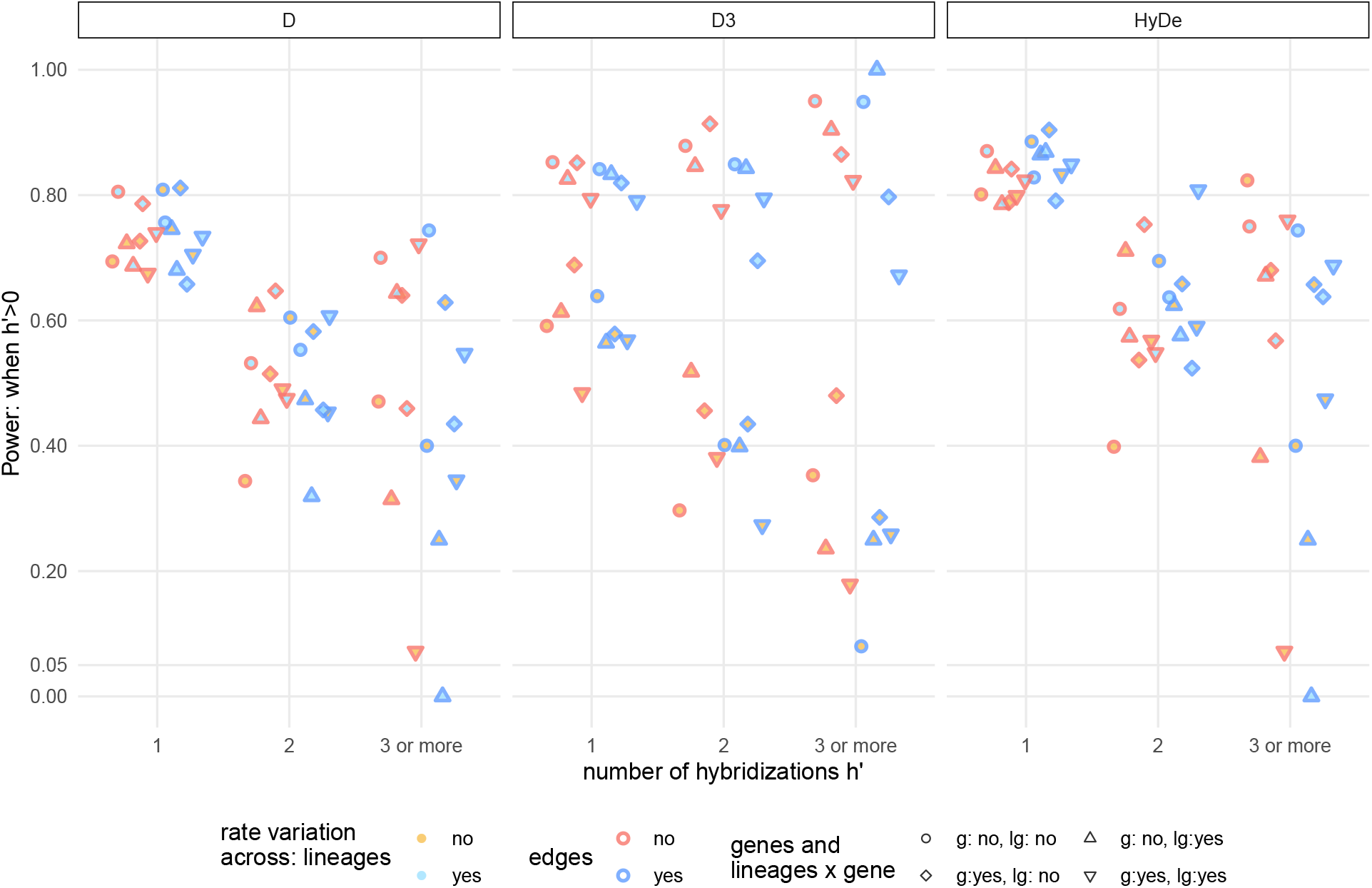
Power to detect the presence of reticulations for *D, D*_3_, and HyDe, subsetted by the number of hybridizations in a replicate’s network after iteratively shrinking 2- and 3-cycles. Each point represents the proportion of tests with a p-value below 0.05 across four-taxon subnetworks sharing the same number of hybridizations (*h*′) from 100 simulated networks, with an average of 659 four-taxon subnetworks per point when *h*′ = 1, 190 subnetworks per point when *h*′ = 2, and 48 subnetworks per point (range of 2-116 subnetworks, point with 2 subnetworks is blue filled and outlined upward pointing triangle) when *h*′ ≥ 3. Fill color, outline color and point shape are as in Fig. 4.

When *h*′ = 2, the power to reject the no-reticulation hypothesis discernibly decreased compared to *h*′ = 1, for both *D* (32%-65%) and HyDe (40%-81%). The range of power for *D*_3_ increased (27%-91%). Compared to the case when *h*′ = 1, *D*_3_’s power increased in the presence of lineage-rate variation, and decreased in the absence of lineage-rate variation. When *h*′ ≥ 3, all three tests have a wide range of power. Some of this variation may result from the low sample size of these scenarios (between 2 and 116 triplet subnetworks had *h*′ ≥ 3) and the resulting low accuracy of our power estimates in this case. The trend remained for *D*_3_, with consistently higher power in the presence than in the absence of lineage-rate variation.

When considering *h* instead of *h*′ to define the alternative hypothesis, the power of all methods declined significantly when *h* = 1 (Fig. S1 in the Appendix), as this condition is a mixture of networks with *h*′ = 1 and networks with *h*′ = 0, considered above for the type-1 error. *D*_3_ is particularly under-powered without lineage-rate variation (∼ 20%), and much more powerful with lineage-rate variation (∼ 80%), as seen for the type-1 error.

## 4 Discussion

### 4.1 Substitution rate variation

Recently, more attention has been paid to the assumptions made by introgression tests, and to the possible incorrect interpretations from these tests if their assumptions are broken. Particularly, ghost lineages can lead to erroneous conclusions of which taxa are involved in introgression for the *D*-statistic [Tricou et al., 2022a], *D*_3_ [Tricou et al., 2022b], and HyDe [Bjørner et al., 2022]. However, few studies have evaluated the effect of substitution rate variation on introgression statistics. In a small simulation of 4 taxa and 5000 unlinked sites, Blair and Ané [2020] increased the gene tree branch lengths for two taxa. They found that *D* was more likely to incorrectly infer introgression as rate variation increased. However, to our knowledge, substitution rate variation’s effect on different hybridization summary statistics has not yet been comprehensively evaluated, let alone the different effects of the various kinds of rate variation.

#### 4.1.1 Rate variation across species lineages has a large effect

Of the various types of rate variation, variable rates across species lineages was the most harmful by far, resulting in the highest levels of type-1 error, across all methods we tested. Comparatively, edge, gene, and lineage-by-gene rate variation had less harmful effects on the type-1 error rate, with rate variation across gene-tree edges having the second most harmful effect.

All three methods were highly susceptible to a violation of their clock assumption, leading to significantly elevated type-1 error when species lineages had different substitution rates. When comparing populations within the same species, or very recently diverged species, it may be assumed that divergence was recent enough to ignore rate variation across populations. Any such assumption should be made very cautiously, given the high sensitivity of *D, D*_3_ and HyDe to this assumption.

The ABBA and BABA patterns used by *D* (and HyDe) occur in low frequency compared to the expected BBAA pattern. They are the result of ILS if the gene tree conflicts with the species tree and if one substitution occurred on the conflicting branch. The presence of a single substitution corresponds to the infinite-site model assumed by the *D* test. However, the ABBA and BABA patterns can also arise from homoplasy on the majority of gene trees that do not conflict with the species tree. Under homoplasy, the derived nucleotide (e.g. B) is gained twice independently, as illutrated in Fig. 7. We can expect homoplasy to have an asymmetric effect on the ABBA and BABA site frequencies if the two sister species evolved at different rates. For example, consider the case when species *t*_2_ evolved faster than its sister *t*_1_ after divergence. Then we can expect more ABBA than BABA sites resulting from homoplasy, in which the same substitution occurred independently in *t*_3_ and one of the two sister species (see Fig. 7), because a substitution in *t*_1_ (BABA) would be less frequent than a substitution in *t*_2_ (ABBA). Under this scenario, the frequency of ABBA and BABA sites caused by ILS would remain the same, as the probability of a conflicting gene tree with one substitution along the internal branch would remain unaffected by rate variation after divergence. Therefore, the sensitivity of *D* and HyDe to rate variation across lineages could, in part, be the result of asymmetric homoplasy, when *D*’s infinite-site assumption is violated.

**Figure 7:**
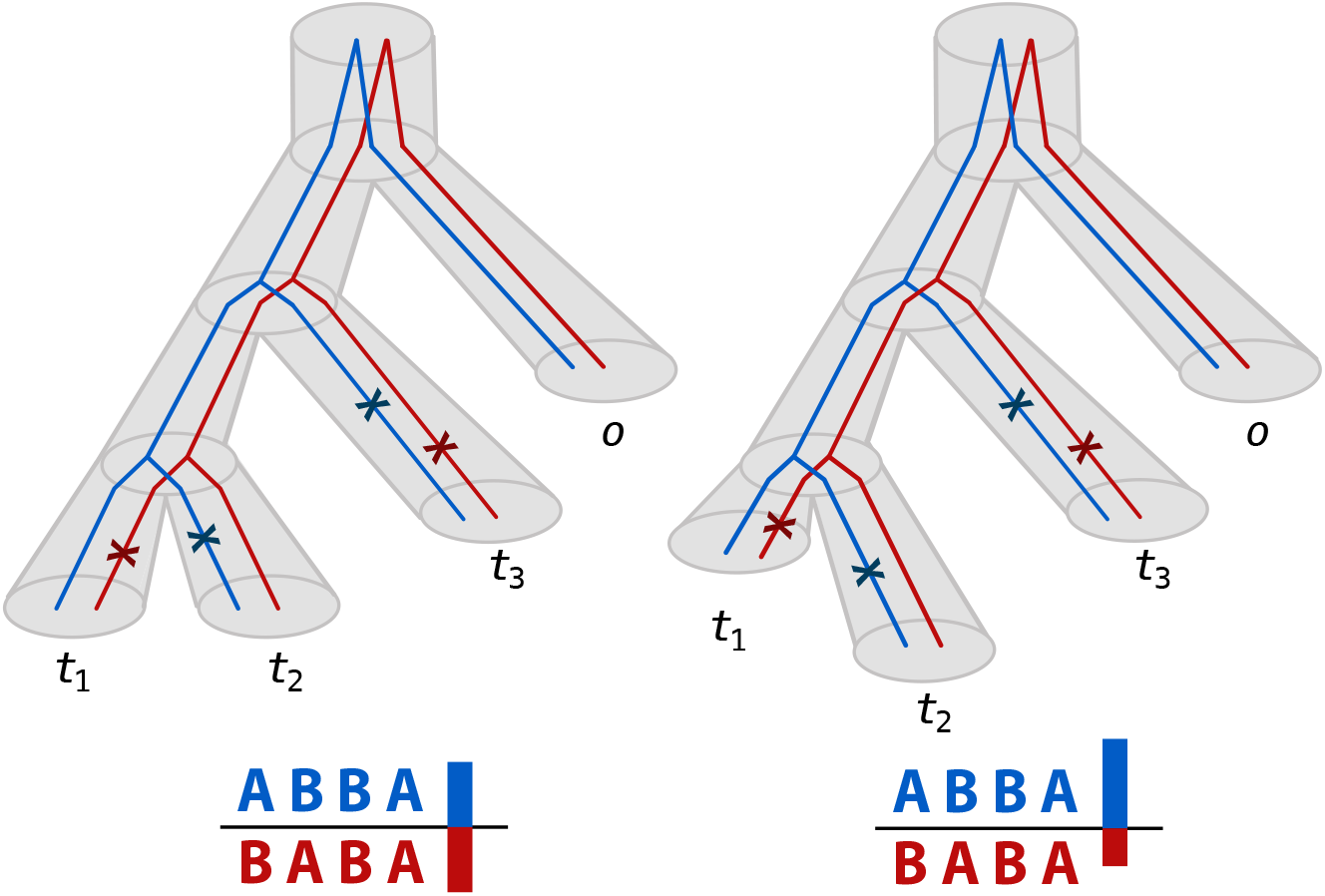
Illustration of the asymmetric effect of homoplasy under a departure from the clock when the two sister species evolved at a different rate, causing elevated type-1 errors for the *D* statistic. Branch lengths are drawn proportional to their number of substitutions per site. Two gene trees (red and blue lines) in agreement with the species tree (gray cylinders) is shown with two homoplastic sites: one with substitutions marked in red and the other in blue. Each site has two independent substitutions, one in the *t*_3_ lineage and the other in the *t*_1_ lineage (resulting in the BABA pattern, in red) or in the *t*_2_ lineage (resulting in the ABBA pattern, in blue). Left: the lineages to *t*_1_ and *t*_2_ have equal numbers of expected substitutions, so *N*_ABBA_ − *N*_BABA_ is expected to be close to 0, and *D* is expected to find no evidence for reticulation. Right: the lineage to *t*_2_ evolved faster than its sister *t*_1_, so homoplasy would result in more ABBA sites than BABA sites, causing the numerator *N*_ABBA_ − *N*_BABA_ of *D* to shift towards positive values, and *D* to lead to a significant test.

#### 4.1.2 *D*_3_ as a test of the molecular clock

By far, *D*_3_ was the most affected by rate variation. The magnitude of its type-1 error rate in our simulations (between 60-80%) differs from that in simulations that did not include rate variation. For example, Bjørner et al. [2022] found that *D*_3_ has a “similar good performance (high precision and low false positive/negative rates) on single shallow hybridizations involving few taxa” compared to HyDe, *D* and other methods (TICR [Stenz et al., 2015], MSCQuartets [Rhodes et al., 2021] and *D*_*p*_ [Hamlin et al., 2020]). *However, Bjørner et al*. *[2022] did not include rate variation in their simulation framework. The power of D*_3_ was moderate to detect reticulations. In contrast, *D*_3_’s error rate (when the true 3-taxon network had no reticulation) and power (when the subnetwork had one or more reticulation) were both greatly increased in the presence of lineage-rate variation. In other words, *D*_3_ was much more sensitive to a departure from the clock assumption, than to a departure from the assumption of no reticulation.

Intuitively, we can expect *D*_3_ to be especially sensitive to rate variation between the two species lineages that are sister. If sister species *t*_1_ and *t*_2_ evolved at different rates after diverging from each other, then we can expect their genetic distance from an outgroup, *d*_12_ and *d*_13_, to differ from each other (Fig. 8). In turn, *D*_3_’s numerator *d*_13_ − *d*_12_ is expected to differ from 0, not because of reticulate evolution, but due to *t*_1_ and *t*_2_ evolving at different rates.

**Figure 8:**
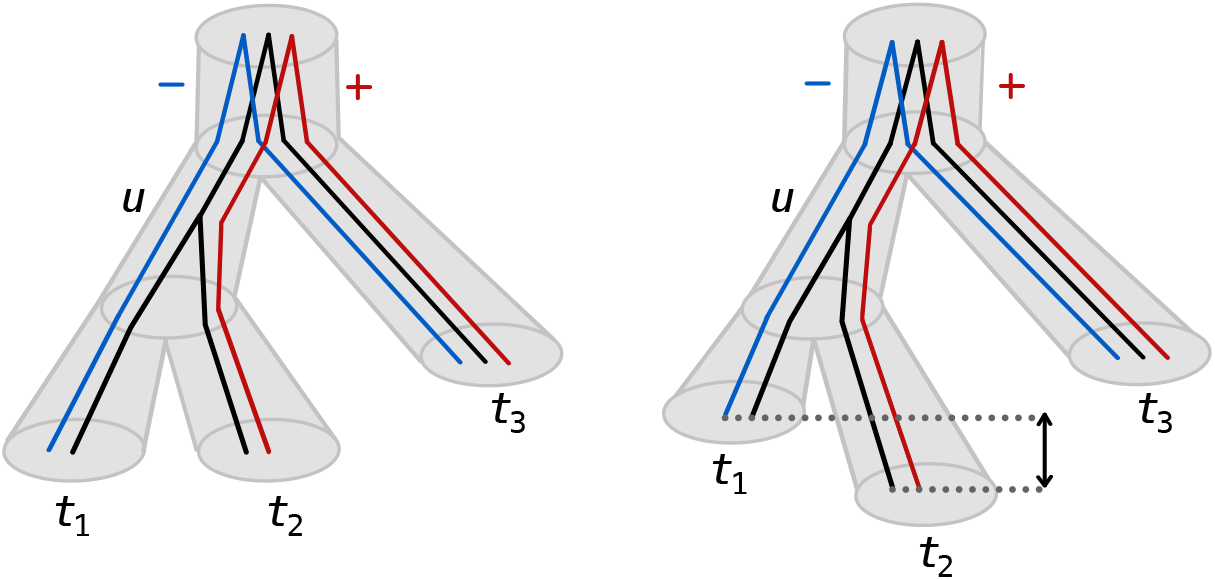
Illustration of the departure from a clock between the two sister lineages *t*_1_ and *t*_2_ on the *D*_3_ statistic. The species tree is shown with gray cylinders. The numerator for *D*_3_ is the difference between *d*_23_ (in red) and *d*_13_ (in blue), with a large absolute value interpreted as evidence for introgression [Hahn and Hibbins, 2019]. Branches in gene trees (thin lines) are color-coded by their contribution to *D*_3_. Branches ancestral to the coalescent event *u* between *t*_1_ and *t*_2_ cancel out. The branch from *u* to *a* contributes negatively, and the branch from *u* and *b* contributes positively. Left: the population lineages to *t*_1_ and to *t*_2_ have equal lengths in substitutions per site, so *d*_23_ − *d*_13_ is expected to be close to 0, and *D*_3_ is expected to find no evidence for reticulation. Right: the lineages to *t*_1_ and to *t*_2_ have different lengths in substitutions per site, so *d*_23_ − *d*_13_ is expected to be far from 0 (shown by the arrow) and *D*_3_ is expected to lead to a significant test.

Compared to the simulation by Tricou et al. [2022b] who did not include rate variation, we considered the true network to include all reticulations in the subnetwork of the 3 taxa used by the *D*_3_ statistic, even if reticulations involved “ghost” lineages who left no sampled descendants among the 3 taxa of interest. That way, type-1 errors in our study correspond to rejecting networks that truly had no reticulations. Going beyond the cautionary message in Tricou et al. [2022b] that any significant *D*_3_ statistics should be interpreted as “the result of an introgression event originating from outside the tree formed by the 3 taxa considered”, we add that any significant *D*_3_ statistics may indicate a departure from a molecular clock. Only if there is ground to assume a lack of rate variation across lineage, then a significant *D*_3_ statistic may be interpreted as a signal of introgression, possibly involving ghost lineages.

#### 4.1.3 HyDe has an elevated type-1 error

Surprisingly, HyDe showed an elevated rate of falsely detecting reticulation, well above 5%, in the ideal case when there was no rate variation. This elevated error rate may be due to a violation of HyDe’s assumption that the hybrid edges have a length of 0, which corresponds to hybrid speciation, or lineage-generative reticulation, and when each parental population has a descendant representative in the 3 ingroup taxa used by HyDe. In our simulations, reticulations were simulated to be lineage-generative (as assumed by HyDe) with probability 0.5, and otherwise lineage-neutral or lineage-degenerative with probability 0.25 each. Both latter types would result in one or both hybrid edges with positive lengths, whereas HyDe assumes zero-length hybrid edges. In addition, subsampling taxa (as done when considering all triplets of ingroup taxa) may result in subnetworks with hybrid edges of non-zero length, again violating HyDe’s assumption. Ji et al. [2022] found HyDe to have an elevated false positive rate, of magnitude similar to that in our study (between 7-13%). Like in our study, their simulations considered a framework that violated HyDe’s assumption of zero-length hybrid edges, with directional introgression between two lineages. In practice, we lack a priori knowledge of the reticulation type and whether HyDe’s assumption hold, so caution should be used when interpreting a positive result by HyDe.

#### 4.1.4 Practical consequences

Our study points to the advantages of methods that do not require assumptions on evolutionary rates across lineages or across genes, including methods based solely on topological information. Indeed, methods that use gene tree topologies (without edge lengths) should be robust to rate variation, to the extent that the inference of gene trees is not affected by long-branch attraction or other topological biases [Molloy and Warnow, 2018]. For the task of estimating species trees (without reticulation), methods based on topological information are known to be more robust than methods that use edge length information from gene trees, due to rate variation between lineages [Liu et al., 2009] or other errors in branch length estimation [DeGiorgio and Degnan, 2014, Degnan, 2018]. While a thorough comparison of network inference methods has not been carried out yet, our study points to a similar conclusion.

Estimating the tree topology of loci requires loci that are long enough, however, with multiple linked SNPs contributing to the estimation of their locus tree. With a handful of SNPs per locus, sequence-based and SNP-based methods like *D, D*_3_ and HyDe are attractive, in part because they don’t require to partition the alignment into loci and to estimate locus trees. When information about linkage between sites is lacking, rate variation could be hard to account for. Desirable alternatives to *D, D*_3_ and HyDe would be new sequence-based methods capable of distinguishing and accounting for different types of rate variation, including rate variation across sites and across species lineages, when there is also variation of the tree topology across sites due to ILS and reticulation.

### 4.2 Hybridization complexity

For methods that aim to infer the full phylogenetic network from the full taxon set, identifiability of the network is not guaranteed if the network is not a tree [Solís-Lemus and Ané, 2016, Baños, 2019, Allman et al., 2022, Xu and Ané, 2023, Allman et al., 2023]. These studies found that most reticulations (those that form large cycles) can be identified by most methods, provided that the network is of “level-1”, that is, reticulations are isolated from one another and form cycles that do not share any edges. These studies also highlighted the difficulty to identify reticulations when they overlap, that is, in networks of level 2 or higher. From methods that only use a small subset of taxa, like *D, D*_3_ and HyDe, we can expect reticulations to have an even stronger lack of identifiability. To date, no theoretical studies and few simulation studies have addressed the identifiability of individual reticulation events by *D, D*_3_ or HyDe, and how this identifiability is affected by the presence of other hybridization events or by the complexity (level) of the network with more than a few reticulations. Often studies limit their simulations only to a single introgression event in a network, not exploring the effect of multiple events [e.g. Tricou et al., 2022a,b].

Our findings align with these theoretical studies of what network inference methods can identify. Notably, we find that 2-cycles and 3-cycles are undetectable by *D, D*_3_ and HyDe. We also find a decrease in the power to detect reticulations when the true number of reticulations increases, across all methods (except *D*_3_ under rate-lineage variation, as *D*_3_ has high power to detect a departure from the clock). In our study, reticulations in the 3-ingroup subnetworks must overlap with each other after shrinking 2/3-cycles, that is, they must form a complex “blob” [Xu and Ané, 2023, Allman et al., 2023]. From theory, we expect multiple reticulations to be difficult to identify. Indeed, the decrease in power that we find shows a tendency for these reticulations to *hide each other*: With an increased number of reticulations in the same group, the site pattern probabilities used by *D, D*_3_ and HyDe tend to look more like site patterns on a tree. This finding is also in line with that of Bjørner et al. [2022], who recently evaluated the performance of a suite of summary statistic hybrid detection methods on a fixed set of networks. They found a high false negative rate (low power) for all tested methods on a 25-taxon and 5-hybridization network which they hypothesized could be due to “a weakening of the hybridization signal when multiple hybridizations are affecting the same taxa.” This tendency for reticulations to hide each other, when they all occur in the same group, calls for methods that can extract richer information from the genomic or multi-locus alignment than *D, D*_3_ and HyDe do, and methods that can infer networks of level higher than 1. More theory is needed, to determine the identifiability (or lack thereof) of reticulations in networks with level 2 or higher, depending on the input data and model assumptions.

## 6 Acknowledgements

We thank Sarah Friedrich for advice and help with figures.

## 7 Supplementary Material

The code to reproduce all analyses and figures is available at https://github.com/cecileane/simulation-ratevariation. Simulated scaled networks and the results from *D, D*_3_, and HyDe available at the Dryad Digital Repository: https://doi.org/10.5061/dryad.qbzkh18nd.

**Figure S1:**
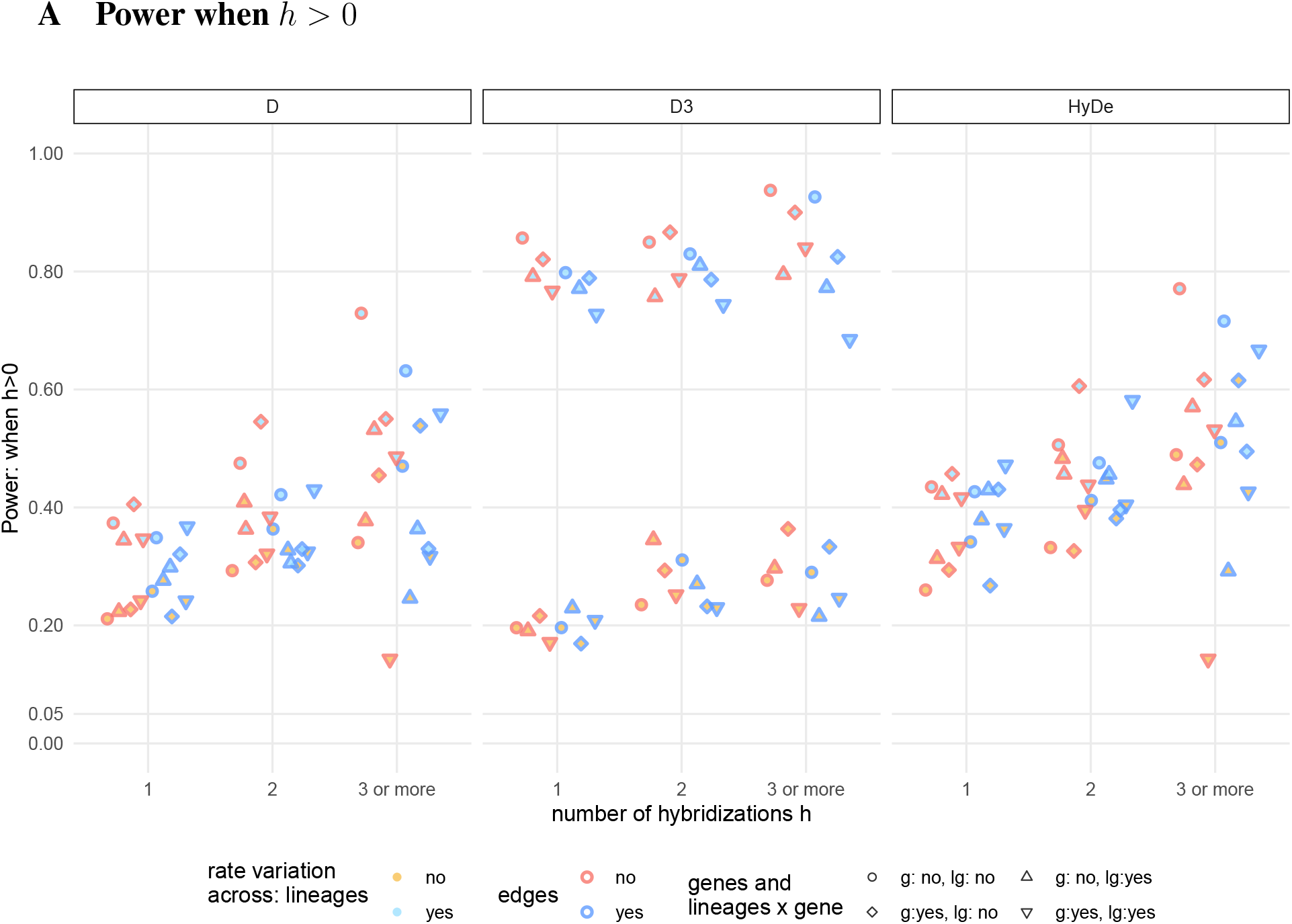
Power to detect the presence of reticulations for *D, D*_3_, and HyDe, depending on the number of hybridizations *h* in the 3-taxon ingroup network before iteratively shrinking 2- and 3-cycles. Each point represents the proportion of tests with a p-value below 0.05 across four-taxon subnetworks sharing the same number of hybridizations in the ingroup (*h*) from 100 simulated networks, with an average of 2193 four-taxon subnetworks per point when *h* = 1, 469 subnetworks per point when *h* = 2, and 100 subnetworks per point when *h* ≥ 3. Fill color, outline color and point shape are as in Fig. 4.

